# Host cell metabolism contributes to delayed-death kinetics of apicoplast inhibitors in *Toxoplasma gondii*

**DOI:** 10.1101/384123

**Authors:** Katherine Amberg-Johnson, Ellen Yeh

## Abstract

*Toxoplasma gondii* and related human parasites contain an essential plastid organelle called the apicoplast. Clinically-used antibiotics and other inhibitors that disrupt apicoplast biogenesis cause a mysterious “delayed-death” phenotype, in which parasite growth is unaffected during the first lytic cycle of inhibitor treatment but is severely inhibited in the second lytic cycle even after drug removal. Critical to understanding the complex downstream cellular effects of these drug classes is the timing of apicoplast loss during inhibitor treatment and how it relates to this peculiar growth phenotype. Here we show that, upon treatment with diverse classes of apicoplast inhibitors, newly-replicated *T. gondii* parasites in the first lytic cycle initially form apicoplasts with defects in protein import or genome replication and eventually fail to inherit the apicoplast altogether. Despite the accumulation of parasites with defective or missing apicoplasts, growth is unaffected during the first lytic cycle, as previously observed. Strikingly, concomitant inhibition of host cell isoprenoid biosynthesis results in growth inhibition in the first lytic cycle and unmasks the apicoplast defects. These results suggest that defects in and even complete loss of the apicoplast in *T. gondii* are partially rescued by scavenging of host cell metabolites leading to death that is delayed. Our findings uncover host cell interactions that can alleviate apicoplast inhibition and highlight key differences in “delayed-death” inhibitors between *T. gondii* and *Plasmodium falciparum*.

## Introduction

The Apicomplexan phylum contains human and animal parasites including *Toxoplasma gondii,* which cause opportunistic infections, and *Plasmodium* spp., which cause malaria. These parasites contain an essential plastid organelle called the apicoplast that is derived from secondary endosymbiosis of a red alga (1–3). While the apicoplast is no longer photosynthetic, it houses essential pathways for biosynthesis of fatty acids, heme, iron-sulfur clusters, and isoprenoid precursors (4–6). Apicoplast ribosome inhibitors, such as clindamycin and doxycycline, are used clinically for treatment of acute toxoplasmosis and malaria chemoprophylaxis, respectively (7, 8). In both *T. gondii* tachyzoites and blood-stage *P. falciparum*, these inhibitors cause a peculiar “delayed-death” phenotype *in vitro*: Treatment with inhibitors during the first lytic cycle has no effect on parasite replication, egress of daughter parasites from the first host cell, or reinfection of new host cells. However, parasites subsequently fail to replicate in the second lytic cycle, even if the inhibitor is removed (9–11). In addition to structurally-diverse antibiotics targeting the prokaryotic ribosome, inhibitors of DNA gyrase (ciprofloxacin) and the apicoplast membrane metalloprotease FtsH1 (actinonin) also cause delayed death in *T. gondii*, indicating that drug properties do not account for the delayed growth inhibition and that delayed death is likely a result of complex downstream cellular effects of apicoplast inhibitors (9, 10, 12, 13).

Each of these apicoplast inhibitors causes defects in apicoplast biogenesis—its growth, division, and inheritance, leading to the formation of *T. gondii* parasites that are missing the apicoplast entirely (10, 12, 13). It is therefore surprising that these drug-treated parasites replicate to wild-type levels in the first lytic cycle during inhibitor treatment, as defects in or loss of the apicoplast should render parasites unable to produce essential apicoplast-derived metabolites (14). How parasites are able to compensate for this loss during the first lytic cycle remains poorly understood. Of note, growth kinetics resembling delayed death have also been observed for inhibitors that block apicoplast metabolic function and genetic disruption of proteins required for apicoplast biogenesis or metabolism, suggesting that inhibiting production of essential apicoplast metabolites may be the common perturbation leading to delayed death in *T. gondii* (5, 15–18).

A number of models have been proposed to explain how apicoplast defects lead to delayed death. One model proposes that apicoplast metabolites are only required for the successful establishment of a parasitophorous vacuole (PV) but are dispensable during intravacuolar replication (9). Another model proposes that growth of parasites with defective apicoplasts during the first lytic cycle is supported by sister parasites with functioning apicoplasts in the same vacuole (19). These models, however, are inconsistent with experiments in which clindamycin-treated parasites were manually released from the host cell prior to completion of the first lytic cycle, separated from sister parasites, and allowed to establish a new infection. These drug-treated, prematurely-lysed parasites were able to establish a new PV and replicate, albeit at reduced rates that depended on the duration of drug treatment and number of replications in the previous vacuole (9). These parasites also eventually fail to replicate in the third or, with continued manual release, later lytic cycles (9), suggesting that the “delay” in growth inhibition is not strictly tied to lytic cycles. Thus neither of the proposed models is sufficient to explain the delayed-death phenotype.

Several key questions remain. First, what is the timing of apicoplast biogenesis defects and loss upon treatment with apicoplast inhibitors? Apicoplast loss is an important downstream cellular consequence of these inhibitors but has not been quantified during a full lytic cycle. Second, do apicoplast inhibitors with distinct molecular targets lead to different rates of apicoplast loss? While the literature would suggest similar phenotypes between diverse classes of apicoplast inhibitors, this has yet to be confirmed with a side-by-side comparison. Third, what is the role of the host cell in delayed death? We hypothesize that, since *T. gondii* replicates in a metabolically active host cell, host metabolites may compensate for apicoplast inhibition. Fourth, how do the downstream cellular effects of apicoplast inhibition differ between *T. gondii* and *P. falciparum?* Their distinct replication cycles, differences in their host cell metabolic activity, and the overlapping but distinct inhibitor classes that cause delayed death in these parasites suggest different mechanisms underlie the seemingly similar delayed growth kinetics.

To address these questions, we validated an apicoplast marker to monitor apicoplast biogenesis defects and loss and show that multiple classes of apicoplast inhibitors cause gradual accumulation of parasites with disrupted or missing apicoplasts. Interestingly, the delayed-death growth kinetics caused by these apicoplast inhibitors is modulated by an inhibitor of host cell isoprenoid biosynthesis. These results clarify the complex downstream cellular effects of apicoplast inhibition in *T. gondii* and their similarities and differences to that in *P. falciparum*.

## Results

### Apicoplast inhibitors cause reduced or absent FNR-RFP, an apicoplast marker

We selected three inhibitors that are well-documented to cause delayed-death in *T. gondii* and have strong evidence for their target in the apicoplast: actinonin (membrane metalloprotease FtsH1), clindamycin (ribosome), and ciprofloxacin (DNA gyrase) (5, 10, 12, 13, 20). During treatment with these compounds, the apicoplast had been previously observed by microscopy of *T. gondii* RH parasites expressing an apicoplast-targeted ferredoxin NADP+ reductase fused to red fluorescence protein (FNR-RFP) (12, 18, 21–24). In these experiments, treatment with apicoplast inhibitors leads to vacuoles with some parasites missing FNR-RFP fluorescence (10, 12, 18). While these studies suggested that the inhibitors lead to disruption of apicoplast biogenesis, it was unclear how exactly FNR-RFP fluorescence corresponded to apicoplast presence. Furthermore, since these studies used microscopy, they could only count apicoplasts in vacuoles containing <8 parasites, corresponding to no more than 3 replications out of >6 total replications during the lytic cycle (10, 12, 18). To clarify these initial observations, we sought to develop a quantitative method to monitor apicoplast loss through a full lytic cycle. Because microscopy of large vacuoles with 8+ parasites is difficult, we used flow cytometry to count and quantify the FNR-RFP fluorescence of individual extracellular parasites after host cell egress.

We compared FNR-RFP fluorescence with the presence of the apicoplast genome, a known apicoplast marker. Briefly, parasites were treated with actinonin, clindamycin, or ciprofloxacin for a single lytic cycle. After natural egress from the first lytic cycle, egressed parasites were collected and their FNR-RFP fluorescence was quantified by flow cytometry (Fig 1A-B). While untreated parasites retained high levels of FNR-RFP fluorescence, treatment with each apicoplast inhibitor generated two populations of parasites: one devoid of FNR-RFP fluorescence [FNR-RFP(-)] and one in which FNR-RFP was detectable but with a mean fluorescence intensity 25-75% less than that of the untreated population [FNR-RFP(reduced)]. We sorted these two populations and quantified their apicoplast:nuclear genome ratio by qPCR (Fig 1C). The apicoplast genome was at reduced levels compared to untreated parasites but still detectable in FNR-RFP(reduced) parasites. In contrast, the apicoplast genome was below the detection limit in FNR-RFP(-) parasites, consistent with loss of the apicoplast in FNR-RFP(-) parasites. These results validate the use of the FNR-RFP fluorescence as a marker for apicoplast presence. It also suggests that, while the FNR-RFP(reduced) cells still contain the apicoplast, the apicoplast is defective given the reduced levels of FNR-RFP fluorescence, lower apicoplast genome copy number, and failure of both populations to grow in the next lytic cycle (10, 12, 13).

**Figurue 1.**
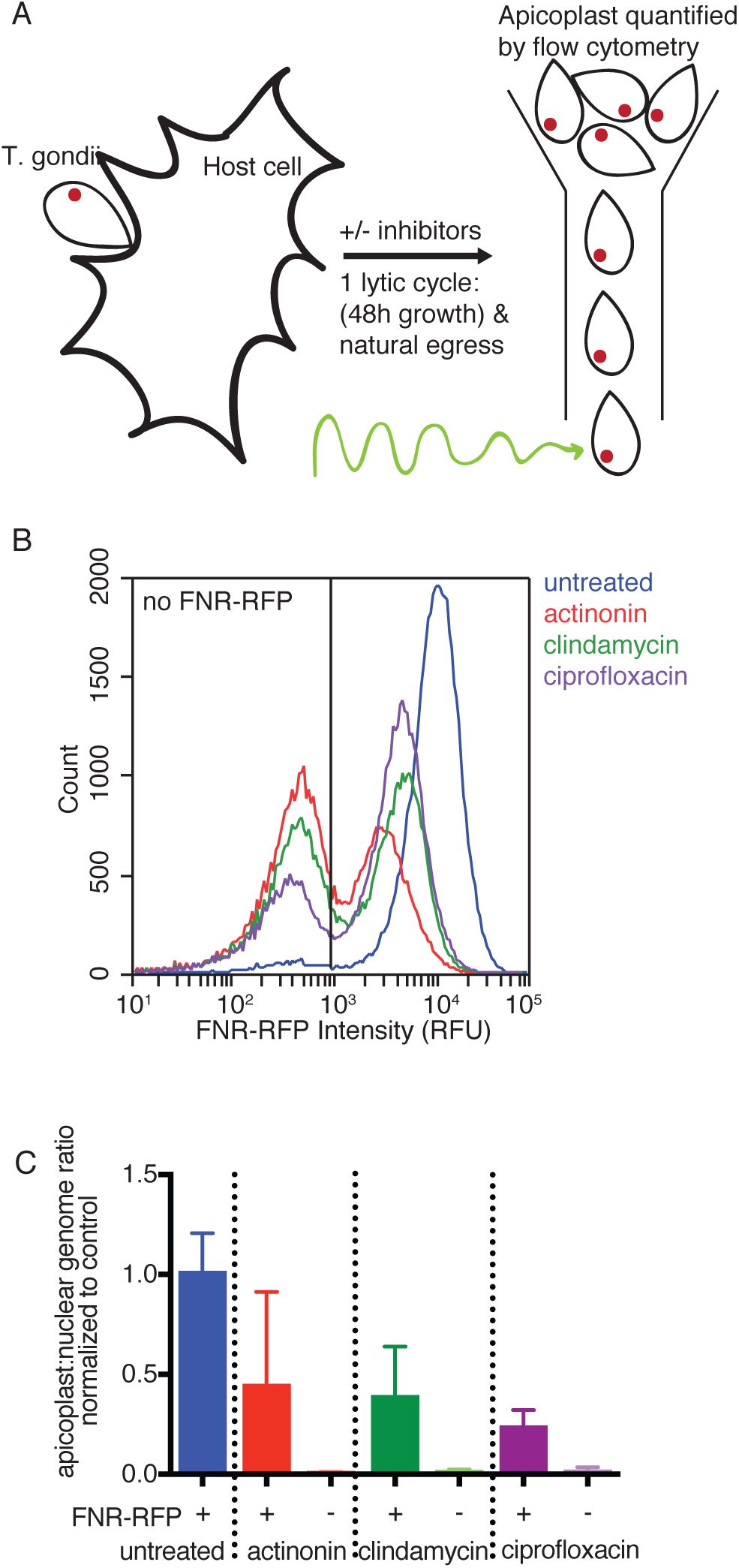
Apicoplast inhibition causes reduced or absent FNR-RFP which is correlated with apicoplast genome levels. (A) Schematic of experimental procedure. *T. gondii* is allowed to infect host cells for a single lytic cycle in the presence or absence of apicoplast inhibitors. Resulting parasites are collected and analyzed by flow cytometry. (B) Representative histogram of FNR-RFP fluorescence intensity of parasites after a single lytic cycle of growth in the presence or absence of apicoplast inhibitors (blue = untreated, red = actinonin, green = clindamycin, purple = ciprofloxacin). Non-fluorescent gate was drawn based off of parasites that did not express FNR-RFP or any other fluorescent marker. (C) Apicoplast:nuclear genome ratio of sorted parasites after a single lytic cycle of growth in the presence or absence of apicoplast inhibitors. Gates to sort FNR-RFP(+) or FNR-RFP(-) parasites were drawn based on parasites that did not express FNR-RFP. Data is representative of two biological replicates performed in technical triplicate. Error bars represent the standard error of the mean (SEM).

### Apicoplast loss occurs gradually during the first lytic cycle

During the *T. gondii* lytic cycle, a single parasite undergoes >6 synchronous rounds of binary division forming >64 daughter parasites within a vacuole in the host cell (25). Apicoplast growth, division, and inheritance is coordinated with parasite replication leading to exactly one apicoplast per parasite (26). Because parasites grow to wild-type levels during the first lytic cycle during treatment, we were surprised to find that the large majority of parasites had either disrupted or undetectable apicoplasts at this time point (Fig 1A-C). We therefore further characterized the timing of apicoplast disruption during the first lytic cycle.

*T. gondii* parasites were treated with apicoplast biogenesis inhibitors and parasites were harvested after 6, 12, 24, 36, or 48 hours of treatment (Fig 2A). At each time point, we manually released parasites from host cells and assessed apicoplast status based on (1) apicoplast genome levels; (2) import of endogenous nuclear-encoded apicoplast proteins (27); (3) the mean FNR-RFP florescence intensity of the population retaining detectable FNR-RFP signal and (4) the percentage of cells containing detectable FNR-RFP (only apicoplast genome levels were assessed at 6h time point).

**Figurue 2.**
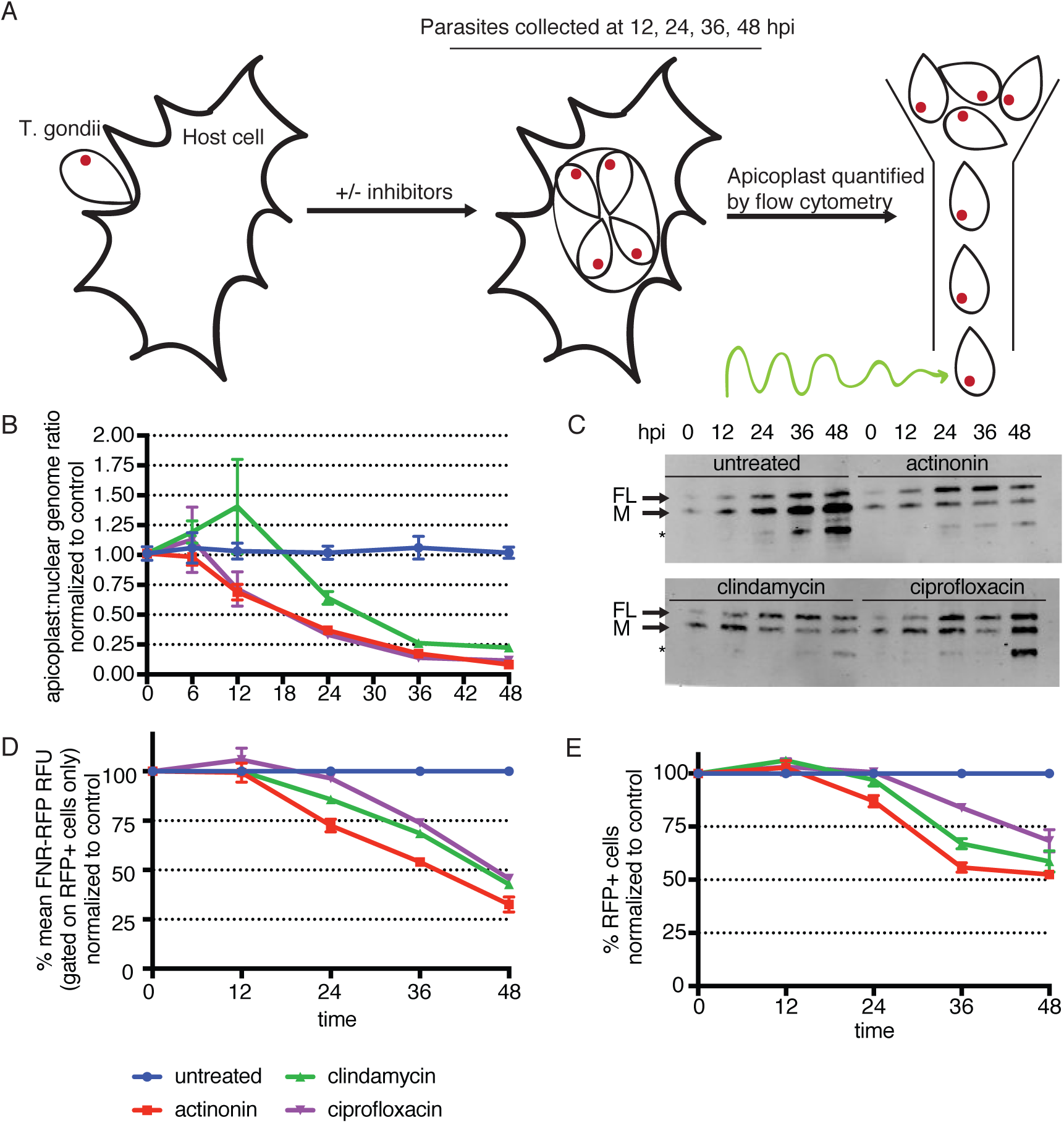
Apicoplast loss occurs gradually over the first lytic cycle of treatment. (A) Schematic of experimental procedure. *T. gondii* parasites are allowed to infect host cells and grow in the presence or absence of apicoplast inhibitors (blue = untreated, red = actinonin, green = clindamycin, purple = ciprofloxacin). At 6, 12, 24, 36, or 48 hours, parasites are manually released from host cells. Because host cell lysis occurs between 36 and 48 hours, parasites analyzed at these time points may reflect growth in the second lytic cycle. (B) Apicoplast:nuclear genome ratio at all time points. Data is representative of two biological replicates performed in technical triplicate. Error bars represent the standard error of the mean (SEM). (C) Western blot of *Tg*Cpn60 from 12-48 hours. FL indicates the full-length protein prior to transit-peptide cleavage. M indicates the mature protein after import into the apicoplast and transit peptide cleavage (27). Data is representative of two biological replicates. (D) The mean FNR-RFP florescence of parasites with detectable FNR-RFP florescence, normalized to control untreated parasites, from 12-48 hours. Data is representative of two biological replicates. Error bars represent the SEM. (E) Percent of cells with detectable FNR-RFP fluorescence, normalized to control untreated parasites, from 12-48 hours. Data is representative of two biological replicates. Error bars represent the standard error of the mean (SEM).

In the first 12 hours of the lytic cycle, *T. gondii* parasites invade host cells, establish a new PV, and divide once or twice. During this time, the only apicoplast defect observed is a slight reduction in the levels of the apicoplast genome for parasites treated with actinonin and ciprofloxacin (Fig 2B). After 24 hours, parasites have completed 3-4 divisions and reductions in levels of the apicoplast genome are detected for all apicoplast inhibitors (Fig 2B). Also at this time point, other apicoplast defects start to appear. We observed that drug-treated parasites begin to accumulate full-length Cpn60 (FL) (Fig 2C), indicating a defect in apicoplast protein import. The FNR-RFP fluorescence of parasites expressing measurable FNR-RFP, starts to dim for parasites treated with actinonin and clindamycin (Fig 2D). Lastly, FNR-RFP(-) parasites start to emerge in samples treated with actinonin (Fig 2E). These defects continue to accumulate for all drug-treated parasites. By 48 hours, *T. gondii* has completed 6-8 divisions and egressed from the host cell. At this time point, the apicoplast genome levels are reduced to 8-22% (Fig 2B), the proportion of (FL) *Tg*Cpn60 has increased (Fig 2C), the mean levels of FNR-RFP fluorescence in the FNR-RFP(reduced) population is 25-50%, and 25-50% of parasites lose FNR-RFP fluorescence all together (FNR-RFP(-)) compared to untreated control parasites. Overall, we observe that *T. gondii* parasites treated with apicoplast inhibitors exhibit apicoplast biogenesis defects as early as the 2^nd^ parasite replication and the severity of these defects worsen throughout the first lytic cycle of treatment, despite normal growth.

### Host isoprenoids are necessary for growth in the first lytic cycle upon treatment with apicoplast inhibitors

Because *T. gondii* parasites treated with apicoplast inhibitors accumulate apicoplast biogenesis defects and lose their apicoplast long before growth inhibition is observed (Fig 2D-E), we sought to determine whether the scavenging of host cell metabolites can compensate for the loss of one or more apicoplast metabolic functions. Isoprenoid biosynthesis is an essential function of the apicoplast (5, 6), and scavenging of host cell isoprenoids by T. gondii has already been shown (28). Therefore parasites were co-treated with apicoplast inhibitors and atorvastatin, a specific inhibitor of host cell isoprenoid biosynthesis. As seen previously, treatment with 13 µM atorvastatin alone did not affect parasite growth, suggesting that in the presence of an intact apicoplast host cell isoprenoid biosynthesis is not essential (Fig 3, Supplemental Fig 1) (28). However, unlike atorvastatin or apicoplast inhibitors alone, the combination of atorvastatin with actinonin, clindamycin, or ciprofloxacin caused parasite growth inhibition within the first lytic cycle (Fig 3, Supplemental Fig 1). Instead of growing to wild-type levels during a full 48-hour lytic cycle, growth defects are detectable starting 24 hours after treatment (Fig 3, Supplemental Fig 1). After 48 hours, parasites co-treated with atorvastatin and apicoplast inhibitors show 30-50% growth compared to parasites treated with apicoplast inhibitors only (Fig 3). The minimum inhibitory concentration (MIC) of apicoplast inhibitors for growth inhibition was similar in the presence and absence of atorvastatin, indicating that atorvastatin’s potential effect on host cell permeability did not result in off-target effects (MIC = 20 µM actinonin +/-atorvastatin; MIC = 4 nM clindamycin +/-atorvastatin). Rather, the altered kinetics of growth inhibition suggest that, with loss of apicoplast function, parasites either require host isoprenoid biosynthesis after 24 hours or deplete host isoprenoid reservoirs after 24 hours. We favor the former model since pretreatment of host cells with atorvastatin for 24 hours prior to infection with *T. gondii* showed similar results (Supplementary Figure 2).

**Figurue 3.**
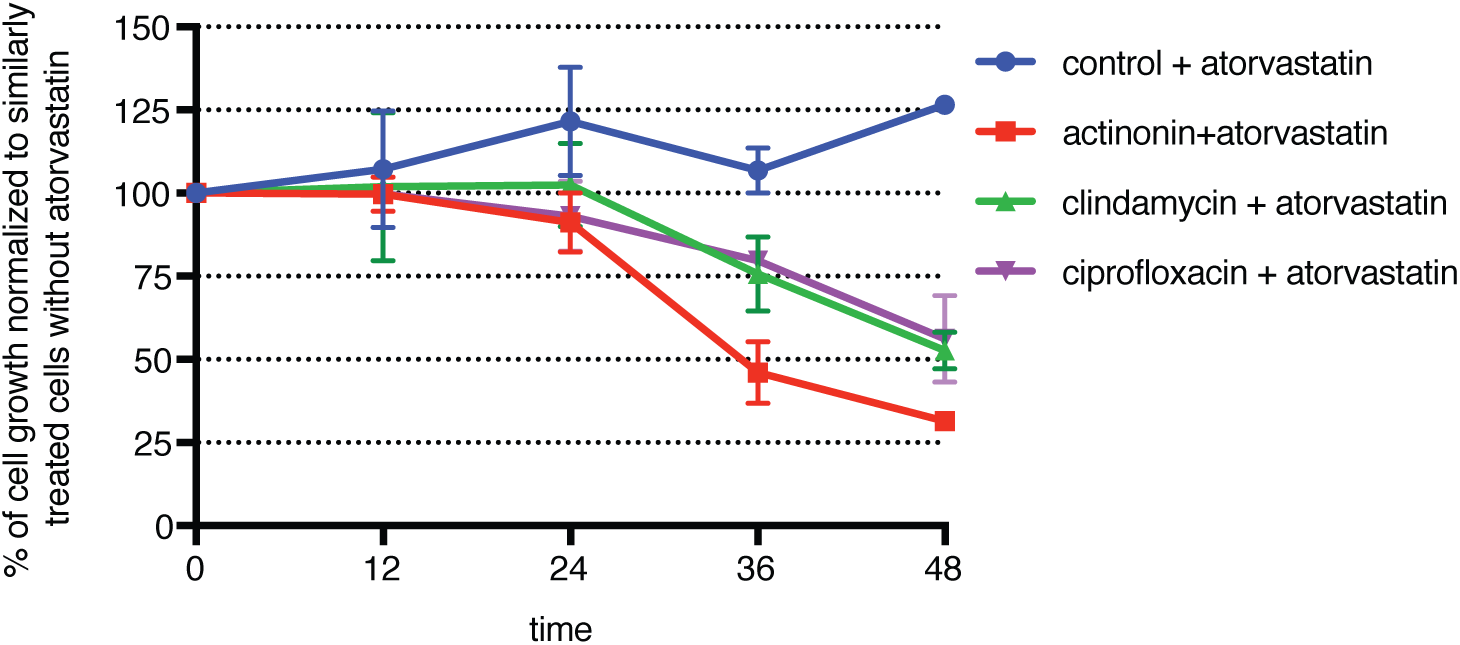
Inhibition of host isoprenoid biosynthesis with atorvastatin results in growth kinetics that deviate from delayed death. (A) Parasite growth quantified by flow cytometry of *T. gondii* manually released from host cells at each time point after treatment with atorvastatin and in the presence or absence of apicoplast drugs (blue = untreated, red = actinonin, green = clindamycin, purple = ciprofloxacin). Growth is normalized to that of parasites treated with the same apicoplast inhibitor but in the absence of atorvastatin at each time point. Data is representative of two biological replicates. Error bars represent the standard error of the mean (SEM). Because host cell lysis occurs between 36 and 48 hours, parasites analyzed at these time points may reflect growth in the second lytic cycle.

## Discussion

Our findings suggest the following model for the downstream cellular effects of apicoplast inhibitors in *T. gondii*. First, several classes of apicoplast inhibitors cause accumulation of apicoplast biogenesis defects and eventual apicoplast loss. We detect these defects beginning as early as the 2^nd^ parasite replication in the first lytic cycle, preceding growth defects in the second lytic cycle.

Second, apicoplast biogenesis defects disrupt its biosynthetic functions important both for intravacuolar growth and likely establishment of infection in new host cells. This model predicts that direct inhibition of apicoplast biosynthetic pathways, without disrupting its biogenesis, will also cause delayed death, consistent with prior observations in the literature (16, 17).

Third, scavenging of host cell metabolites in the first lytic cycle can substitute for metabolites normally biosynthesized in the apicoplast. We specifically tested the scavenging of host cell isoprenoids, which was required 24 hours into the first lytic cycle. Host fatty acids and heme may also compensate for loss of these apicoplast biosynthetic functions, although we were unable to directly test these pathways given the lack of specific inhibitors. Overall, we propose that host cell metabolites support the growth of *T. gondii* during the first lytic cycle of apicoplast inhibition. Our results add to a growing body of work indicating that access to host metabolites regulates the essentiality of parasite metabolic pathways and the organelles that provide them (28–32).

Finally, host cell metabolites cannot compensate for the apicoplast indefinitely, since parasites treated with apicoplast inhibitors ultimately fail to replicate in later lytic cycles. It is possible that metabolites sourced from the apicoplast are required at the beginning of the lytic cycle for PV formation and host cell remodeling. If this is true, then drug-treated parasites prematurely lysed from host cells may continue to grow in subsequent lytic cycles because the apicoplast is still partially functional at these early time points. Alternatively, a combination of host cell scavenging and accrued apicoplast metabolites may be sufficient during the first lytic cell but ultimately become depleted in subsequent lytic cycles. More experiments are required to differentiate between these scenarios, and ultimately reveal the essential products of the *T. gondii* apicoplast.

These downstream cellular effects of apicoplast biogenesis inhibitors in *T. gondii* differ from the known effects in *P. falciparum* in significant ways. First, we show that *T. gondii* parasites lacking apicoplasts accumulate over multiple parasite replications during the first lytic cycle. In contrast, blood-stage *P. falciparum* only undergoes a single round of parasite replication during each host lytic cycle, and apicoplast biogenesis is largely unaffected by treatment with apicoplast translation inhibitors during the first lytic cycle (a small reduction in apicoplast genome copies is sometimes observed) (11, 33). Second, we show that *T. gondii* parasites lacking apicoplasts are viable during the first lytic cycle, and apicoplast loss and parasite growth inhibition are temporally separated. In contrast, *P. falciparum* growth inhibition occurs concurrently with apicoplast biogenesis defects in the second lytic cycle most likely because apicoplast loss cannot be overcome by scavenging of host cell metabolites (6, 11). We suspect this is due to the different metabolic activities of their respective host cells: while *P. falciparum* grows in relatively inert red blood cells, *T. gondii* makes its home in metabolically-active host cells. Third, while all known apicoplast inhibitors cause delayed death in *T. gondii*, only the subset of apicoplast inhibitors that disrupt apicoplast genome expression cause delayed death in *P. falciparum* (12). The common defect leading to delayed death in *T. gondii* appears to be loss of apicoplast metabolic function, while the common defect in *P. falciparum* delayed death is loss of apicoplast genome expression. While it is possible that different molecular targets account for these different inhibitor phenotypes, all apicoplast inhibitors used in this study have strong evidence for the same target in both organisms (10, 12, 13, 33). Instead, these different downstream cellular effects of apicoplast inhibitors likely reflect the different biology of the parasites and their dependence on apicoplast metabolic function.

## Materials and methods

### Chemicals

Actinonin was purchased from Sigma Aldrich and 25 mM aliquots were prepared in ethanol. Clindamycin was purchased from Sigma Aldrich and 5 µM aliquots were prepared in water. Ciprofloxacin was purchased from Sigma Aldrich and 50 µM aliquots were prepared in water. Atorvastatin was a gift from the Smolke lab at Stanford and 50 µM aliquots were prepared in DMSO.

### *Toxoplasma gondii* culture

*T. gondii* RH FNR-RFP (26) was a gift from Boris Striepen (University of Pennsylvania). Parasites were maintained by passage through confluent monolayers of human foreskin fibroblasts (HFFs) host cells. HFFs were cultured in DMEM (Invitrogen) supplemented with 10% FBS (Fetal Plex Animal Serum from Gemini, West Sacramento, CA), 2 mM L-glutamine (Gemini), and 100 µg penicillin and 100 µg streptomycin per mL (Gibco Life Technologies), maintained at 37 C and 5% CO_2_. Parasites were harvested for assays by syringe lysis of infected HFF monolayers.

### Growth inhibition assays

1.5 million extracellular tachyzoites were counted by flow cytometry and allowed to infect T25 flasks containing confluent human foreskin fibroblasts (HFFs). This amount of parasites was chosen because it leads to lysis of the host monolayer after a 48-hour lytic cycle. Infected cells were then incubated with either no apicoplast inhibitor, 40 µM actinonin, 100 nM clindamycin, or 25 µM ciprofloxacin. When included, 13 µM atorvastatin was used. At designated time points during the first lytic cycle, parasites were released from HFFs using syringe lysis and either counted using flow cytometry (BD Accuri C6 Sampler) or collected for qPCR (Applied Biosystem 7900HT) or immunoblot. All time course experiments were repeated with at least 2 biological replicates.

### Flow cytometry and sorting

Fluorescence activated cell sorting (Sony) was performed on FNR-RFP parasites grown for a full lytic cycle in either no drug, 40 µM actinonin, 100 nM clindamycin, or 25 µM ciprofloxacin. Untagged parasites were used to gate on the FNR-RFP(-) cells. One million cells were sorted from each population and frozen down for subsequent analysis.

Flow cytometry (BD Accuri C6 Sampler) was performed to count parasites and quantify FNR-RFP fluorescence. Untagged parasites were used to gate on the FNR-RFP(-) cells. At each time point, syringe lysed cells were washed, resuspended in PBS, and 10 µL fixed volumes were quantified. Samples were always resuspended in PBS directly prior to measurement in the flow cytometer.

### Quantitative real-time PCR

At each time point, syringe lysed parasites (1mL of culture, representing 1/4^th^ of the total sample) was collected, spun down, and frozen prior to analysis. DNA was purified using DNAeasy Blood and Tissue (Qiagen, Germany). Primers were designed to target genes found on the apicoplast or nuclear genome: tufA (apicoplast) 5’-TGGAGCCGCACAAATGGAT-3’/5’-CTTTAGTTTGTGGCATTGGCCC-3’ and actin (nuclear) 5’-GCGCGACATCAAGGAGAAGC-3’/5’-CATCGGGCAATTCATAGGAC-3’ (34). Reactions contained template DNA, 0.15 µM of each primer, and 1x SYBR Green I Master mix (Roche). qPCR reactions were performed at 56C primer annealing and 65C template extension for 35 cycles on a Applied Biosystem 7900HT system. Relative quantification of target genes was determined (35). For each time point, the apicoplast:nuclear genome ratio was calculated relative to the appropriate control collected at the same time. The apicoplast:nuclear genome ratio was measured by qPCR two times.

### Immunoblot

Syringe lysed parasites (1mL of culture, representing 1/4^th^ of total sample) were washed with PBS and frozen down in 1x NuPAGE LDS Sample Buffer (Invitrogen) prior to analysis. Proteins were separated by electrophoresis on 4–12% Bis-Tris gel (Invitrogen) and transferred to a nitrocellulose membrane. After blocking, membranes were probed with 1:5000 polyclonal rabbit anti-TgCpn60 (gift from Boris Striepen, University of Pennsylvania) and 1:10,000 IRDye 800RD donkey anti-rabbit (LiCor Bioscience, Lincoln, NE). Fluorescence antibody-bound proteins were detected with Odyssey Imager (LiCor Biosciences). Immunoblots were performed 2 times.

### Statistical Analysis

When applicable, data was analyzed using Graph Pad Prism software and expressed as mean values ± standard error of the mean (SEM). Basic experiments were repeated at least twice including both positive and negative controls. Biological replicates were performed on different days or on independent cultures while technical replicates were performed using cells from the same culture. Experiments were not blinded. All new reagents were validated prior to use. All qPCR primers were assessed for single amplicon.

## Acknowledgements

We thank Dr. Boris Striepen for providing the *T. gondii* FNR-RFP and the *Tg*Cpn60 antibody. This project has been funded with federal funds from the NIH under Award Numbers 1K08AI097239 (EY), 1DP5OD012119 (EY), and T32GM007276 (KAJ). Funding was also provided by the Burroughs-Wellcome Fund (EY), Chan-Zuckerberg Biohub (EY), and the Stanford Bio-X SIGF William and Lynda Steere Fellowship (KAJ). The funders had no role in study design, data collection and interpretation, or the decision to submit the work for publication.

---

**Supplemental Figure 1.**
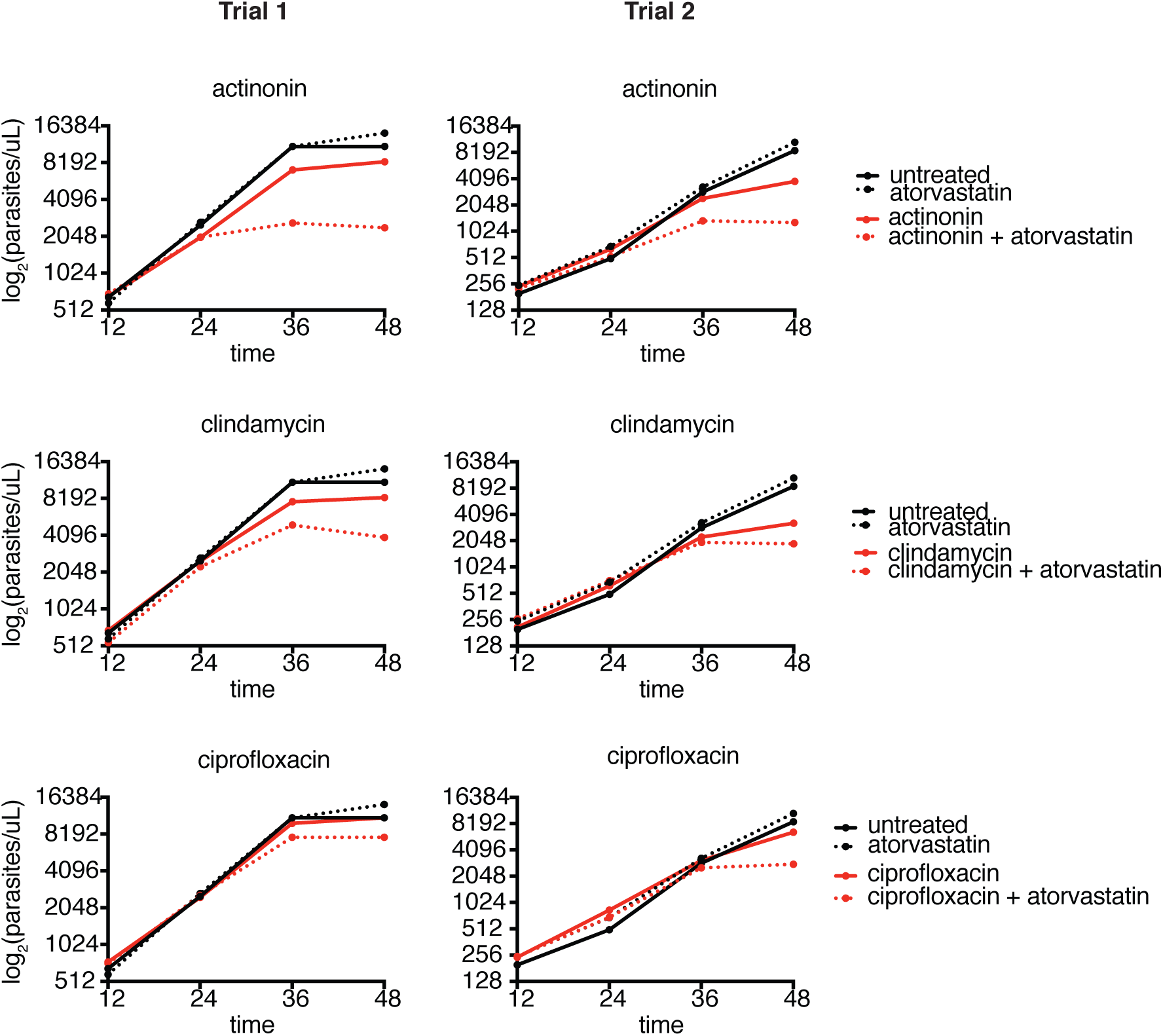
Raw growth curves demonstrate that co-treatment with atorvastatin and apicoplast inhibitors leads to earlier growth defects than treatment with either inhibitor alone. Parasite growth quantified by flow cytometry of *T. gondii* manually released from host cells at each time point after treatment with or without atorvastatin or apicoplast inhibitors. Each biological replicate is plotted separately with the respective controls from that experiment.

**Supplemental Figure 2.**
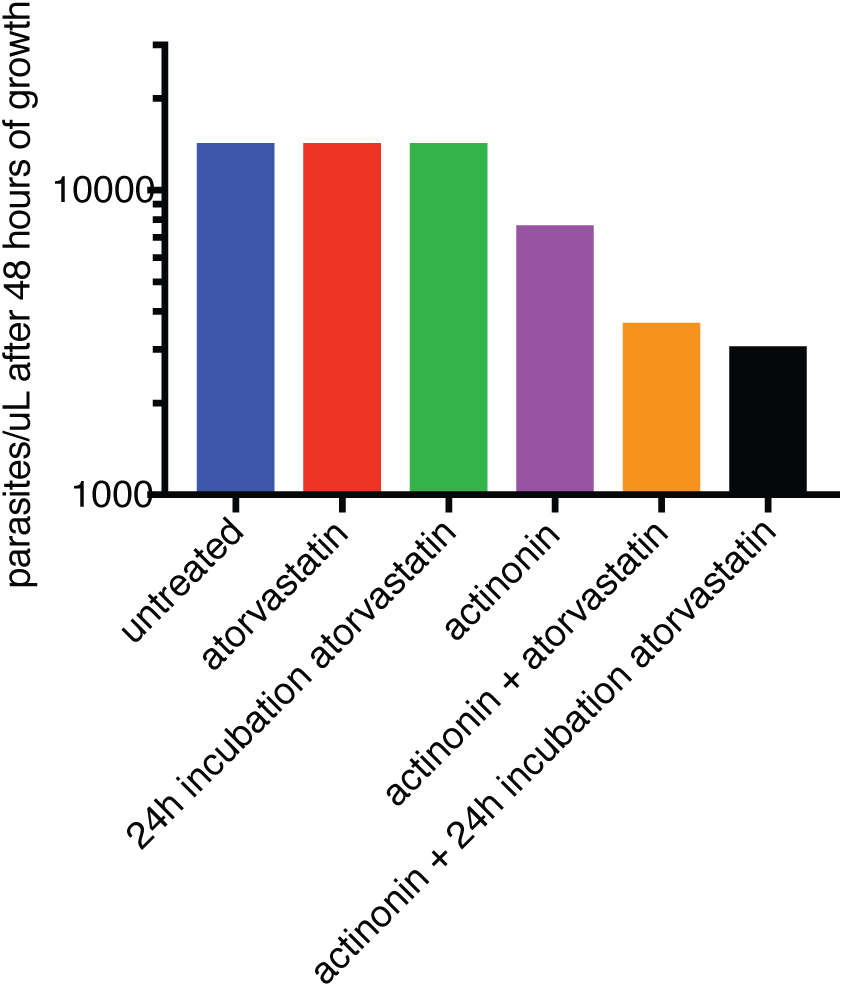
Pre-incubation of host cells with atorvastatin for 24 hours prior to infection does not exacerbate growth defect of cells co-treated with atorvastatin and apicoplast inhibitors. 1.5 million parasites were allowed to infect T25 flasks containing confluent human foreskin fibroblasts (HFFs) that were either untreated or pre-incubated with atorvastatin for 24 hours prior to infection. Infected cells were then incubated with either no inhibitor, atorvastatin only, actinonin only or actinonin+atorvastatin. Parasite growth at the end of a 48 hour lytic cycle was quantified by flow cytometry. Results are from a single biological replicate.

